# Reconstruction and functional annotation of *Ascosphaera apis* full-length transcriptome via PacBio single-molecule long-read sequencing

**DOI:** 10.1101/770040

**Authors:** Dafu Chen, Yu Du, Xiaoxue Fan, Zhiwei Zhu, Haibin Jiang, Jie Wang, Yuanchan Fan, Huazhi Chen, Dingding Zhou, Cuiling Xiong, Yanzhen Zheng, Xijian Xu, Qun Luo, Rui Guo

## Abstract

*Ascosphaera apis* is a widespread fungal pathogen of honeybee larvae that results in chalkbrood disease, leading to heavy losses for the beekeeping industry in China and many other countries. This work was aimed at generating a full-length transcriptome of *A. apis* using PacBio single-molecule real-time (SMRT) sequencing. Here, more than 23.97 Gb of clean reads was generated from long-read sequencing of *A. apis* mecylia, including 464,043 circular consensus sequences (CCS) and 394,142 full-length non-chimeric (FLNC) reads. In total, we identified 174,095 high-confidence transcripts covering 5141 known genes with an average length of 2728 bp. We also discovered 2405 genic loci and 11,623 isoforms that have not been annotated yet within the current reference genome. Additionally, 16,049, 10,682, 4520 and 7253 of the discovered transcripts have annotations in the Non-redundant protein (Nr), Clusters of Eukaryotic Orthologous Groups (KOG), Gene Ontology (GO), and Kyoto Encyclopedia of Genes and Genomes (KEGG) databases. Moreover, 1205 long non-coding RNAs (lncRNAs) were identified, which have less exons, shorter exon and intron lengths, shorter transcript lengths, lower GC percent, lower expression levels, and fewer alternative splicing (AS) evens, compared with protein-coding transcripts. A total of 253 members from 17 transcription factor (TF) families were identified from our transcript datasets. Finally, the expression of *A. apis* isoforms was validated using a molecular approach. Overall, this is the first report of a full-length transcriptome of entomogenous fungi including *A. apis*. Our data offer a comprehensive set of reference transcripts and hence contributes to improving the genome annotation and transcriptomic study of *A. apis*.

## 1. Introduction

Chalkbrood is a widespread disease of the honeybee caused by *Ascosphaera apis* (Maassen ex Claussen) Olive and Spiltoir [1–2], an entomopathogenic fungus that exclusively infects western honeybee larvae. Recently, *A. apis* was reported to infect the larvae of eastern honeybee drones and workers [3]. This brood disease weakens colony productivity and honey production by lowering the number of newly emerged bees and, under certain circumstances, may result in colony losses [4].

The transcriptome can provide the information associated with the number and variety of intracellular genes and uncover the physiological and biochemical processes at a molecular level [5]. To date, an array of technologies has been developed and applied for transcriptome sequencing. Among these, short-read sequencing (i.e., Illumina and Ion Torrent) has become a useful tool for precisely analyzing RNA transcripts and gene expression levels [6–7]. However, most second-generation sequencing (also known as next-generation sequencing (NGS)) platforms offer a read-length shorter than the typical length of a eukaryotic mRNA, including a methylated cap at the 5’ end and poly-A at the 3’ end. To overcome the limitation of short-read sequences, single-molecule real-time (SMRT) sequencing (Pacific Biosciences of California, Inc., CA, USA) was developed, which can produce kilobase-sized sequencing reads, thus eliminating the need for sequence assembly [8–9]. For example, the average read length of PacBio SMRT sequencing is around 10 kb and the subread length can reach up to 35 kb [9]. The full-length transcriptome based on long reads can be used for the exploration and functional characterization of genes, the collection of large-scale long-read transcripts with complete coding sequences, and the identification of gene families [10–11]. However, the technology has a high sequencing-error rate (~15%) when compared to Illumina sequencing (~1%); and it can not currently be directly used to quantify gene expression [12–13]. Fortunately, the limitations of SMRT can be algorithmically improved and corrected by short and high-accuracy sequencing reads [14–15]. Hence, hybrid data derived from SMRT and NGS can offer high-quality and more complete assemblies for genome and transcriptome studies [16–17].

The genome of *A. apis* was published in 2006 with a total size of 20.31 Mb [18]. This version of the reference genome (AAP 1.0) is composed of 8092 contigs which are further assembled into 1627 scaffords [18]; however, it is yet to be fully assembled into complete chromosomes. Transcriptome analysis is a powerful tool for uncovering the relationships between genotypes and phenotypes, leading to a better understanding of the underlying pathways and genetic mechanisms controlling cell growth, development, immune defense, and so forth [19–21]. Our group previously *de novo* assembled and annotated a transcriptome of *A. apis* using short reads from NGS [22]. Based on this reference transcriptome, we further investigated the transcriptomic alteration and pathogeneisis of *A. apis* during the infection process of two different bee species, *Apis mellifera ligustica* and *Apis cerana cerana* [23–24]. To provide a high-quality transcriptome of *A. apis*, in this work, the *A. apis* mycelia were subjected to third-generation sequencing (TGS) using the PacBio Sequel™ system (PacBio, Menlo Park, CA, USA). In parallel, Illumina paired short RNA reads generated separately from *A. apis* mycelia were used to support the SMRT data. Functional annotation of the transcriptome was performed followed by prediction and analysis of long non-coding RNAs (lncRNAs) and transcription factors (TFs). Overall, to the best of our knowledge, this is the first documentation of PacBio-based transcriptomic data of fungi including *A. apis*.

## 2. Results

### 2.1. PacBio SMRT sequencing and error correction of long reads

The workflow of the current work is presented in **Figure 1**. To obtain a representative full-length transcriptome for *A. apis*, the mycelia of *A. apis* were sequenced using PacBio Sequel system, and a total of 13,302,489 subreads (about 23.97 Gb) were yielded from the long-read sequencing, with an average read length of 1802 bp and an N50 of 3077 bp. To provide more accurate sequence information, circular consensus sequences (CCS) were generated from subreads that passed at least once time through the insert, and 464,043 CCS with a mean length of 2970 bp were gained (**Figure 2A**). By detecting the sequences, 402,415 were identified as being full-length (containing a 5’ primer, 3’ primer and the poly-A tail) and 394,142 were identified as being full-length non-chimeric (FLNC) reads with low artificial concatemers (**Figure 2B, Table 1**). The mean length of the FLNC reads was 2820 bp (**Figure 2B, Table 1**). FLNC reads with similar sequences were clustered together using the Iterative Clustering for Error Correction (ICE) algorithm, and 182,165 unpolished consensus isoforms with a mean length of 2701 bp were obtained (**Figure 2C**). In total, 121,776 high-quality isoforms and 58,307 low-quality isoforms were gained after polishing these unpolished consensus isoforms with the Quiver algorithm. Further, the aforementioned low-quality isoforms were corrected using the NGS short reads with Proovread software, resulting in significant improvement in sequence accuracy (**Figure 3**). Finally, we identified 174,095 corrected isoforms with a mean read length of 2728 bp and an N50 length of 3543 bp (**Table 2**).

**Figure 1.**
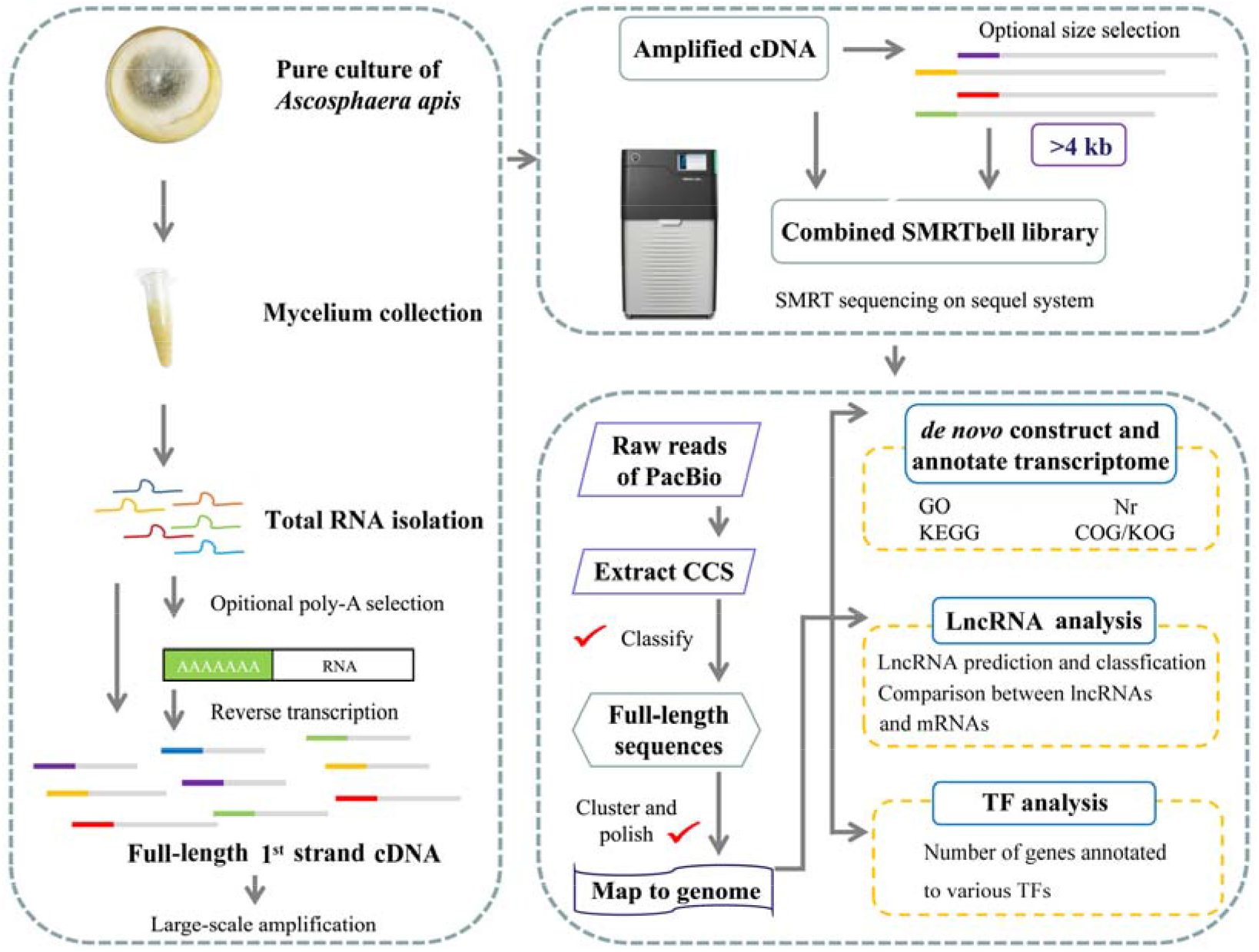
Workflow for fungal mycelia preparation, Iso-seq, data processing and bioinformatic analysis.

**Figure 2.**
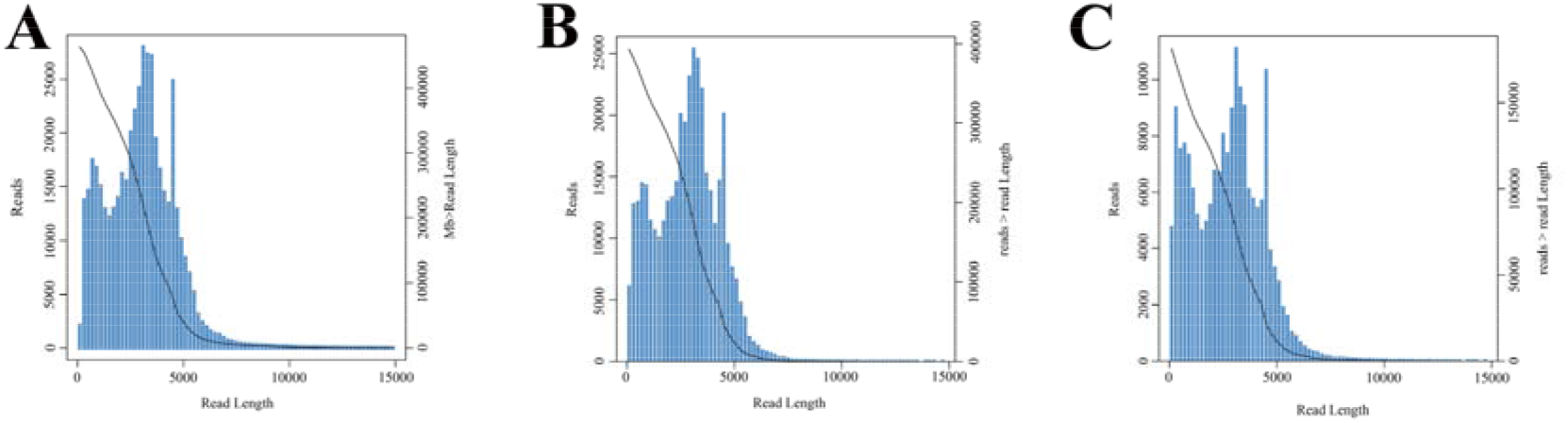
Length distribution of PacBio SMRT sequencing. (A) Number and length distribution of CCS. (B) Number and length distribution of FLNC reads. (C) Number and length distribution of corrected isoforms.

**Figure 3.**
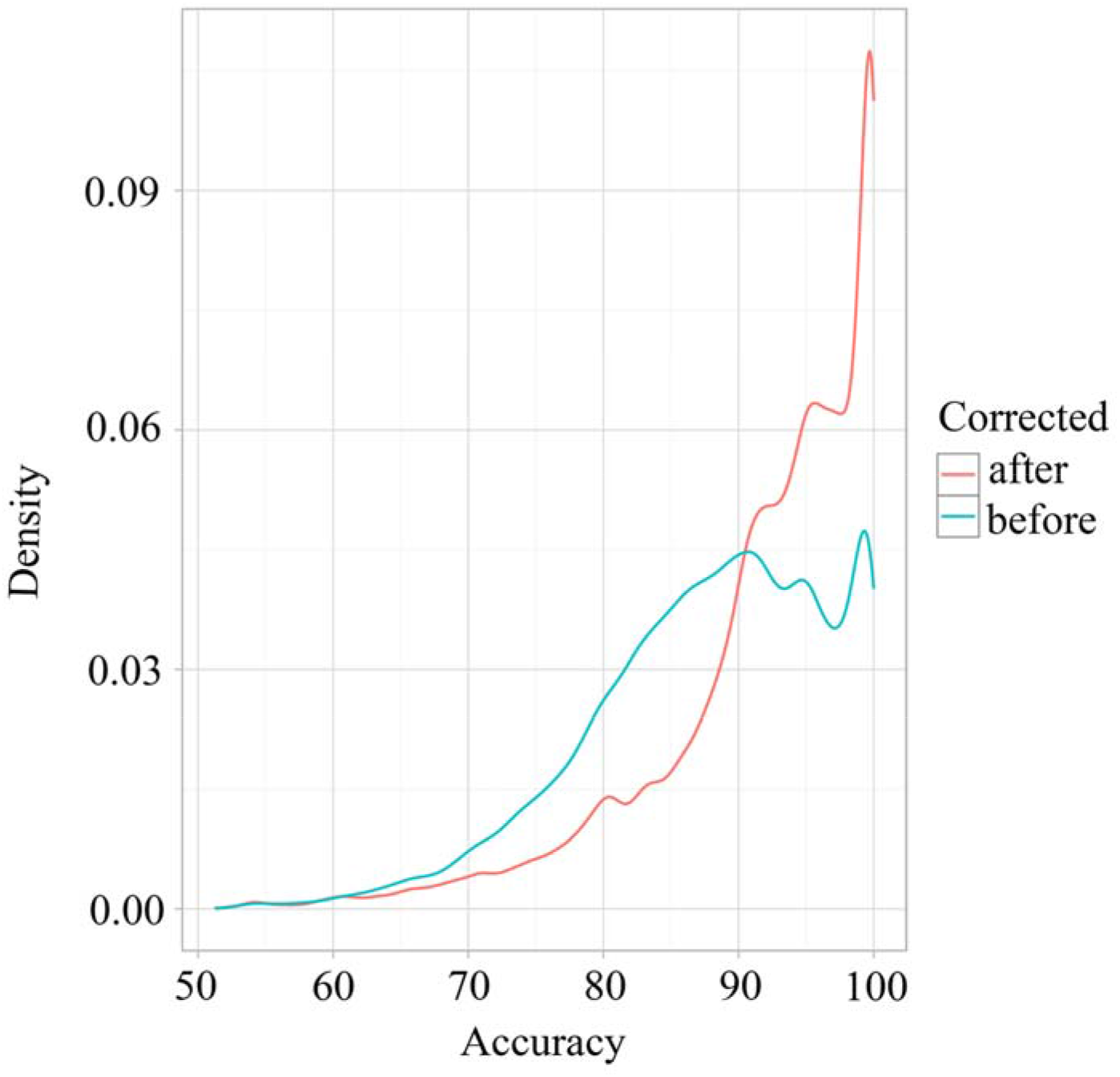
Accuracy of low-quality isoforms after correction with Illumina short reads.

**Table 1.**
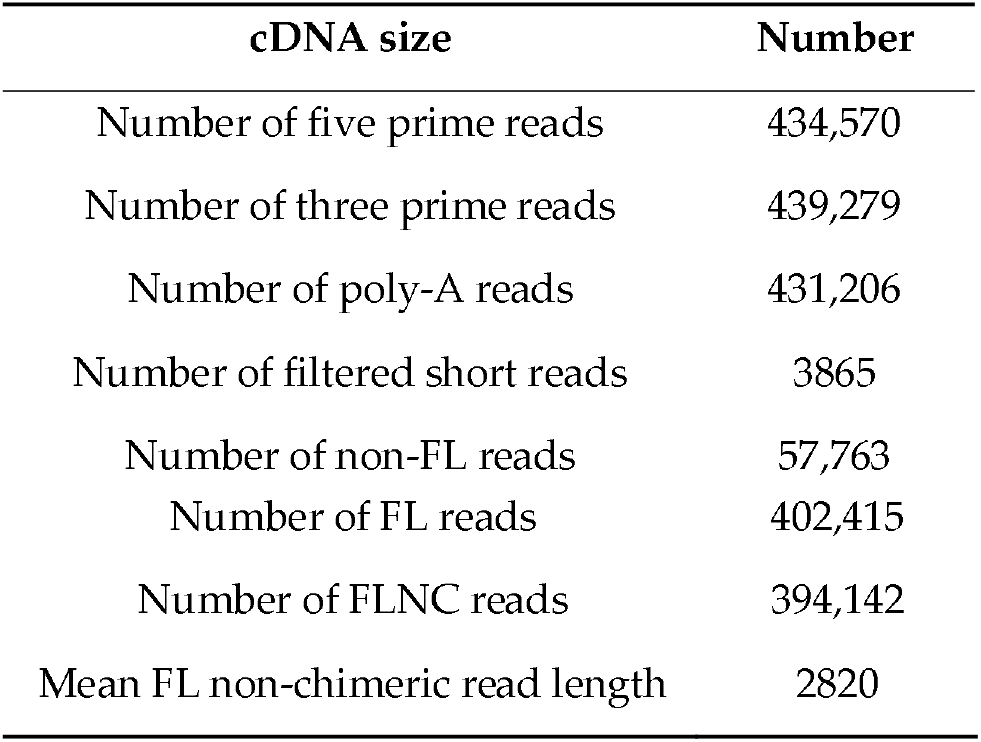
PacBio SMRT sequencing output statistics.

| cDNA size | Number |
| --- | --- |
| Number of five prime reads | 434,570 |
| Number of three prime reads | 439,279 |
| Number of poly-A reads | 431,206 |
| Number of filtered short reads | 3865 |
| Number of non-FL reads | 57,763 |
| Number of FL reads | 402,415 |
| Number of FLNC reads | 394,142 |
| Mean FL non-chimeric read length | 2820 |

**Table 2.** Overview of *A. apis* isoforms after correction.

|  | **Aam** |
| --- | --- |
| Number of total reads | 174,095 |
| Length of total reads (bp) | 474,928,820 |
| Maximum length of total reads (bp) | 13,808 |
| Minimum length of total reads (bp) | 50 |
| Average length of total reads (bp) | 2728 |
| N50 length of total reads (bp) | 3543 |
| GC content of total reads (%) | 49.00 |

### 2.2. Reconstruction of A. apis full-length transcriptome

Here, 84.9% of the FLNC reads were mapped to the *A. apis* genome (**Figure 4A**). In addition, all 174,095 corrected isoforms were aligned against the reference genome of *A. apis*, and 168,740 (96.92%) reads were mapped to the reference genome, including 165,206 (94.89%) unique mapped reads and 3534 (2.03%) multiple mapped reads (**Figure 4B**); 84,734 (48.67%) reads were mapped to the positive strand of the genome, while 80,472 (46.22%) reads were mapped to the opposite strand of the genome (**Figure 4B**). A total of 17,195 isoforms from 5141 genetic loci were mapped to the *A. apis* genome gene set (AAP 1.0), which contains 6442 isoforms from 6442 genetic loci. We identified 3167 known isoforms in reference genome, 2405 novel genic loci from unannotated regions and 11,623 new isoforms from various exons.

**Figure 4.**
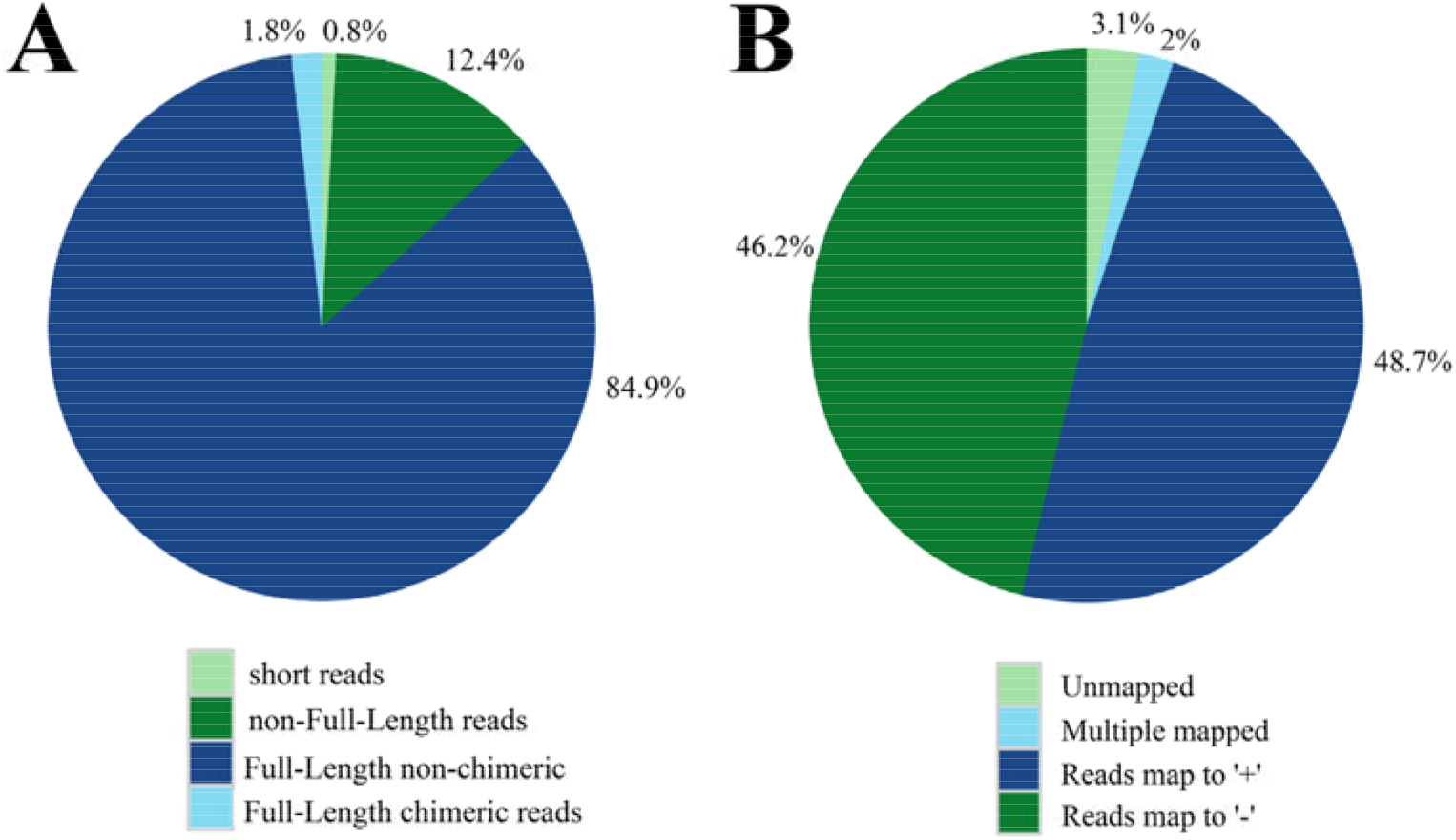
Mapping of SMRT reads (**A**) and corrected isoforms (**B**) to *A. apis* genome.

### 2.3. Functional annotation of the full-length transcriptome of A. apis

Functional annotations of the non-redundant transcripts were determined by searching in the public databases. The results showed that 16,049 (93.34%), 10,682 (62.12%), 4520 (26.29%) and 7253 (42.18%) of the 17,195 isoforms could be found in the NCBI non-redundant protein sequences (Nr), Clusters of euKaryotic Orthologous Groups (KOG), Kyoto Encyclopedia of Genes and Genomes (KEGG) and Gene Ontology (GO) databases, respectively (**Figure 5 A-C**). In addition, the transcripts had the highest number of hits to the *A. apis* (11,441 hits, 88.83%) proteins, followed by *Blastomyces dermatitidis* (118 hits, 0.92%) and *Histoplasma capsulatum* (97 hits, 0.75%) proteins (**Figure 5 D**).

**Figure 5.**
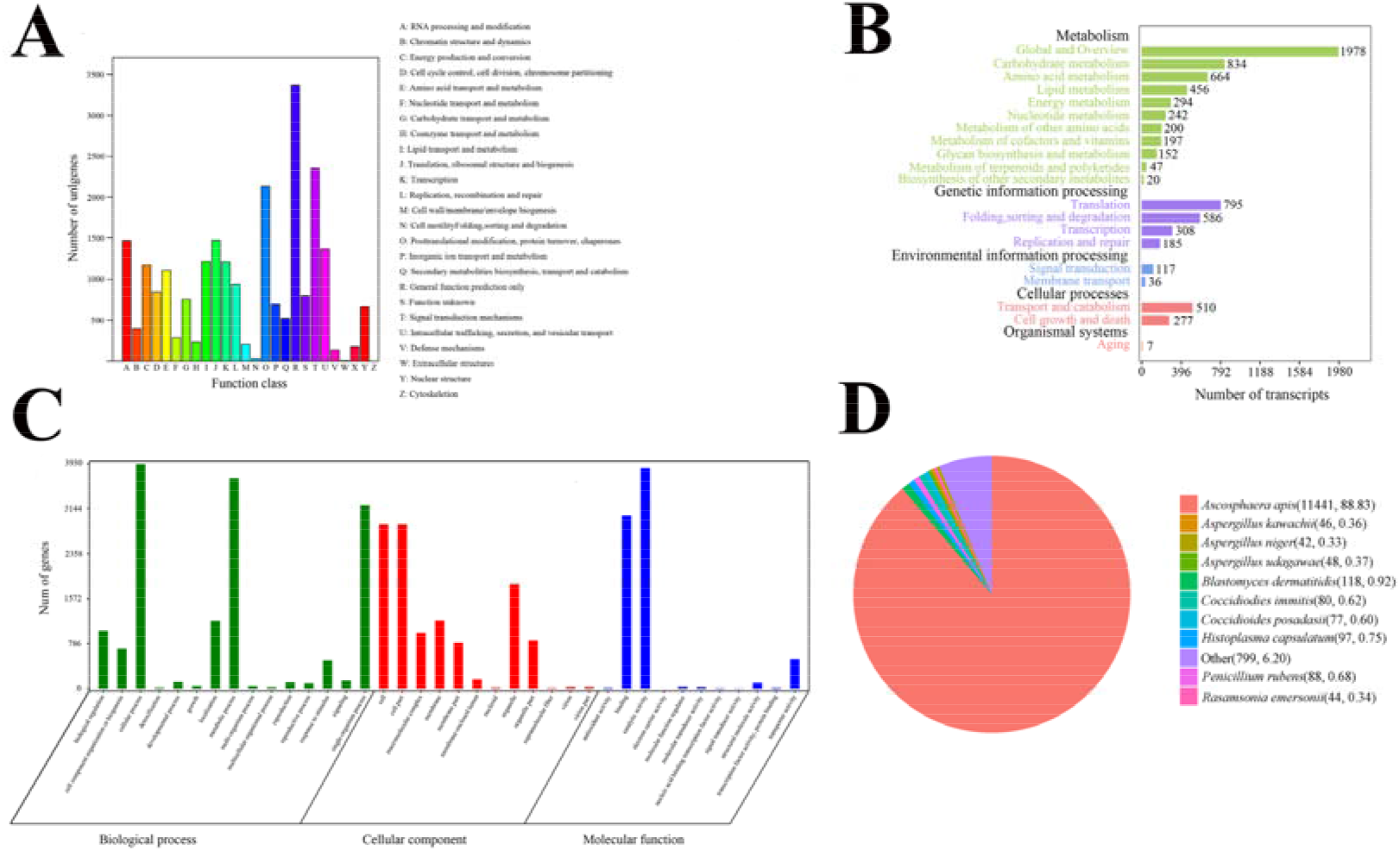
Function annotation of corrected isoforms. (A) KOG classifications of transcripts. (B) KEGG pathways enriched by transcripts. (C) Distribution of GO terms for all annotated transcripts in biological process, cellular component and molecular function. (D) RefSeq Nr Homologous species distribution diagram of transcripts.

### 2.4. LncRNA and TF identification

LncRNAs have been reported to play vital regulatory roles in a wide range of biological processes [25]. The number of lncRNAs predicted by each prediction method is presented in Figure 6A. In total, 1205 high-confidence lncRNAs were identified with an average length of about 912 bp. LncRNAs were classified into five groups according to their biogenesis positions relative to the protein-coding genes of AAP 1.0 annotations: 19.00% (229) of them were generated from intergenic regions, 0.58% (7) from the intronic regions, 15.44% (186) from the sense strand, 18.59% (224) from the antisense strand and 24.65% (297) belong to bidirectional lncRNA (**Figure 6B**). In addition, the majority (72.37%) of the lncRNAs were single exons, and this percentage was obviously higher than that of mRNAs (20.51%) (**Figure 6C**). We additionally observed that when compared with protein-coding transcripts, non-coding transcripts had fewer exons, shorter exon and intron lengths, shorter transcript lengths, lower GC percentages, lower expression levels, and fewer comparison of alternative splicing (AS) events (**Figure 6D-I**), which are similar to findings in other species [26–29].

**Figure 6.**
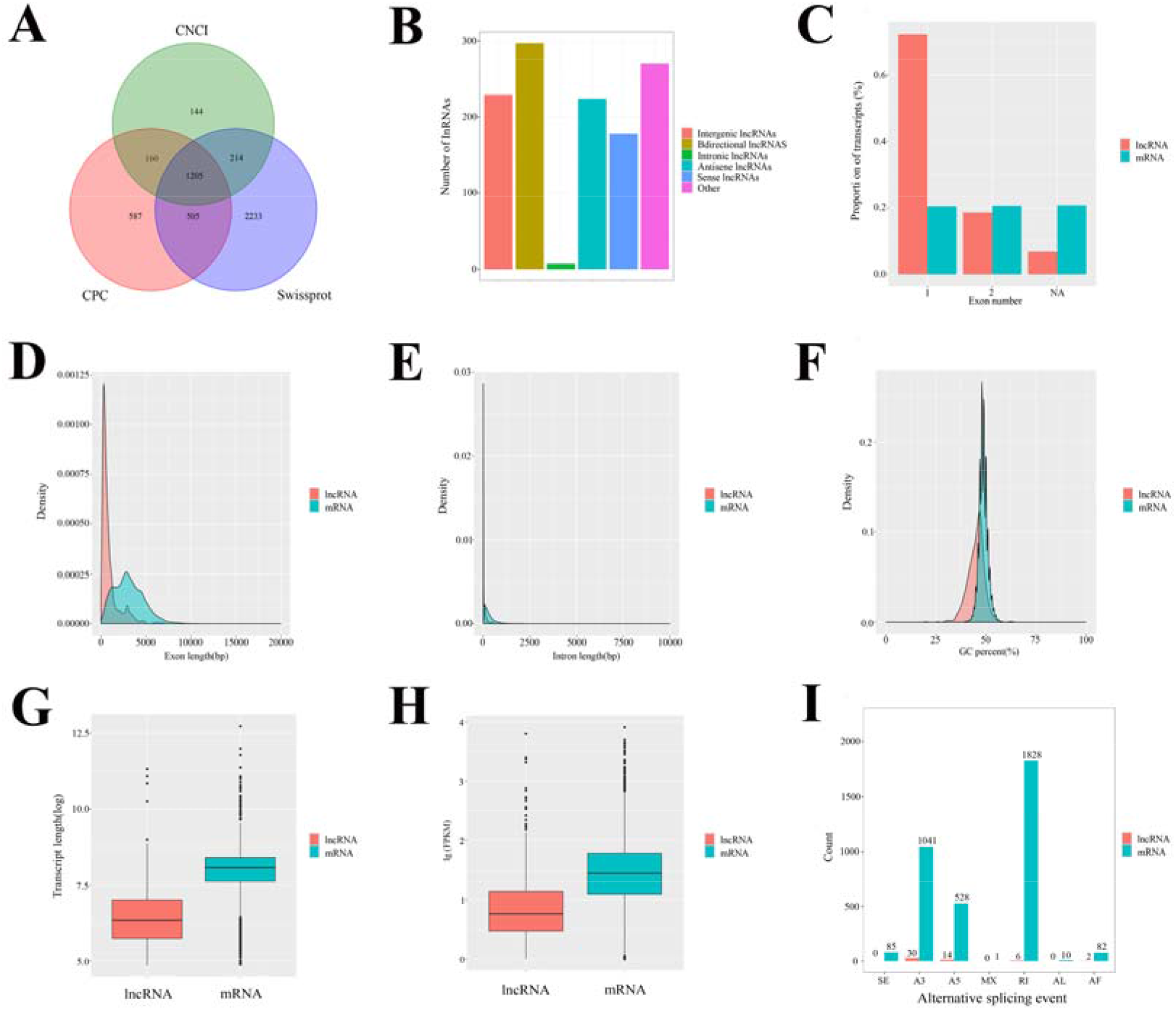
Identification of *A. apis* lncRNAs. (A) Venn diagram of lncRNAs predicted by Coding Potential Calculator (CPC), Coding-Non-Coding Index (CNCI), and pfam methods. (B) Proportions of different types of lncRNAs. (C) Comparison of exon number between lncRNAs and mRNAs. (D) Comparison of exon length between lncRNAs and mRNAs. (E) Comparison of intron length between lncRNAs and mRNAs. (F) Comparison of GC content between lncRNAs and mRNAs. (G) Comparison of transcript length between lncRNAs and mRNAs. (H) Comparison of expression level between lncRNAs and mRNAs. (I) AS events between lncRNAs and mRNAs.

TFs are key components involved in the transcriptional regulatory system in various animals, plants, and insects [12,30]. A total of 253 members from 17 TF families were identified from our transcript datasets. The top 10 TF families were C2H2 (89), bHLH (29), bZIP (29), HB-other (23), HSF (22), MYB_related (19), C3H (13), GATA (8), M-type (5), and NF-YA (3) (**Figure 7**). However, considering the lack of reports associated with *A. apis* TFs, more evidence is needed to confirm our prediction.

**Figure 7.**
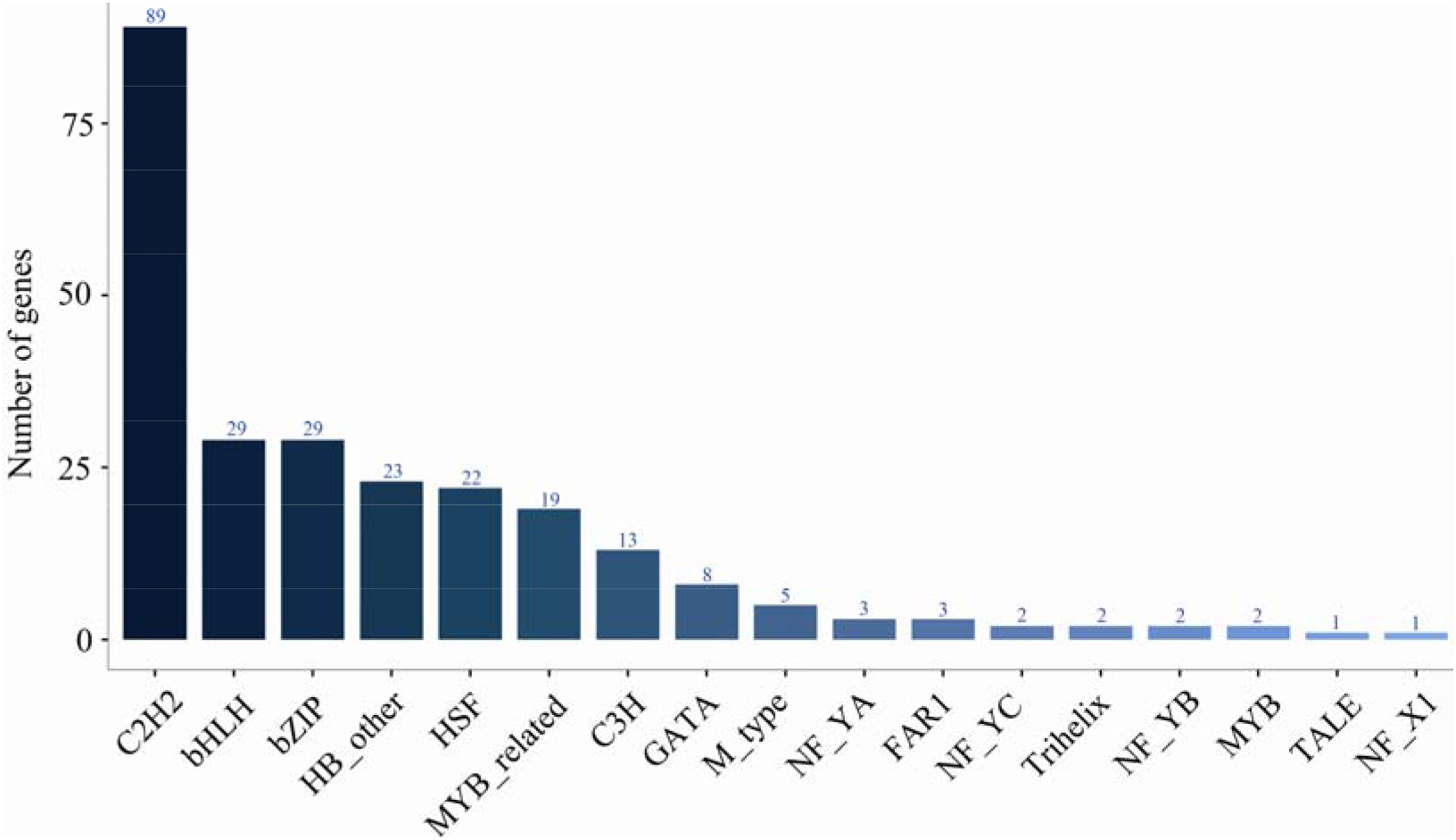
Identification of TFs in *A. apis* mycelia. The number and family of TFs were predicted by SMRT.

### 2.5. Molecular validation of A. apis isoforms

In this work, 16 isoforms were randomly selected for RT-PCR to confirm the expression of novel isoforms. As shown in **Figure 8A**, the signal fragments were successfully detected by agarose gel electrophoresis. In addition, one of these fragments was subjected to molecular cloning and Sanger sequencing, the result further validated the reliability of *A. apis* isoforms (**Figure 8B**).

**Figure 8.**
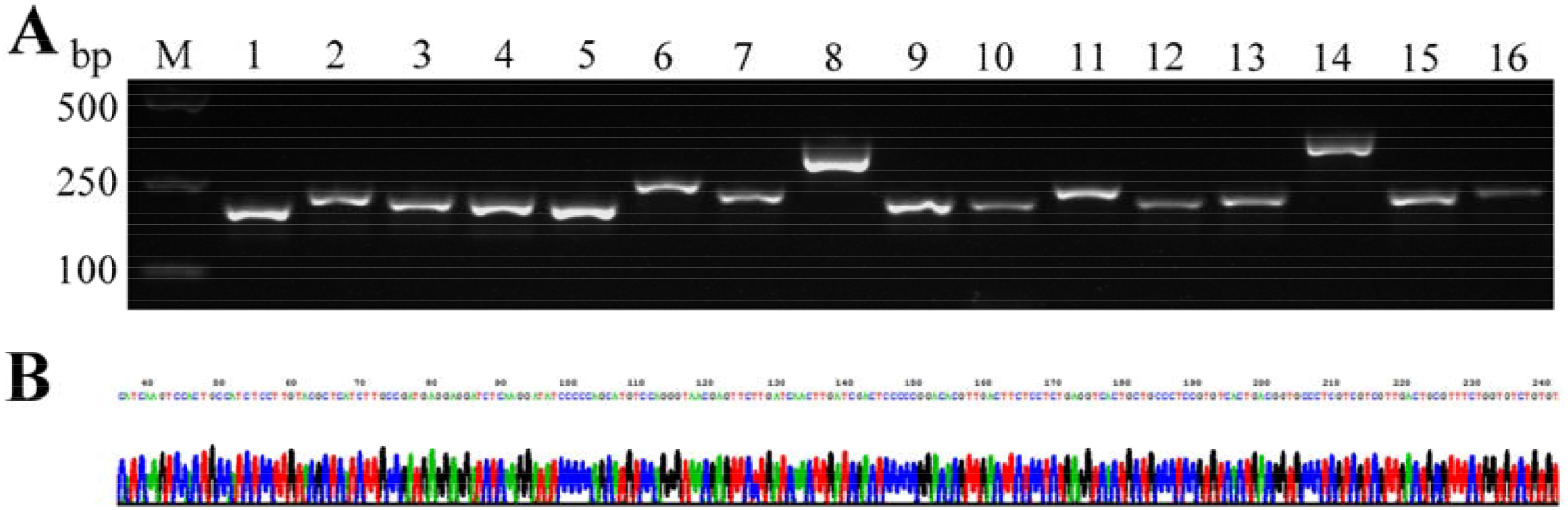
RT-PCR and Sanger sequencing validation of *A. apis* isoforms. (A) Agarose gel electrophoresis of RT-PCR products from 16 *A. apis* isoforms; Lane 1: Isoform000014; Lane 2: Isoform000021; Lane 3: Isoform000027; Lane 4: Isoform000036; Lane 5: Isoform000042; Lane 6: Isoform000085; Lane 7: Isoform000094; Lane 8: Isoform000113; Lane 9: Isoform000127; Lane 10: Isoform000018; Lane 11: Isoform000019; Lane 12: Isoform000028; Lane 13: Isoform000029; Lane 14: Isoform000047; Lane 15: Isoform000063; Lane 16: Isoform000066; Lane M: DNA marker. (B) Sanger sequencing of amplified fragment from Isoform000014.

## 3. Discussion

*A. apis* is a widespread fungal pathogen of the honeybee, but its molecular and omics study is lagging due to a lack of high-quality genome and transcriptome data. Though the genome of *A. apis* was published as early as 2006 [18], the functional annotation was not available until 2016. Transcriptome construction and annotation, particularly for species without a reference genome or a complete genome, has greatly improved with the development and revolution of sequencing techniques and plays a critical role in gene discovery, genomic signature exploration, and genome annotation [31–32]. Our group previously sequenced the *A. apis-infected* honeybee larval guts using the Illumina HiSeq platform, and *de novo* assembled and annotated a transcriptome using the short reads from *A. apis* [22]. Transcriptome analysis is a powerful tool for unraveling the relationships between genotype and phenotype, allowing a better understanding of the underlying pathways and molecular mechanisms regulating metabolism, growth and development, and the immune system [21,33–35]. Based on the previously assembled transcriptome, we conducted a further comprehensive transcriptomic investigation of *A. apis* infecting larvae from the western honeybee and eastern honeybee [23–24]. However, it still remains challenging to reliably assemble full-length from the short reads, and such transcripts are essential to explore post-transcriptional processes, such as AS and alternative polyadenylation (APA) events.

PacBio SMRT sequencing provides better completeness to the sequencing of both the 5’ and 3’ ends of cDNA molecules; thus, it is a superior strategy for the direct generation of a comprehensive transcriptome with precise AS isoforms and novel genes [36–37]. Yi et al. identified 33,300 full-length transcripts (transcript N50 of 5234 bp) of *Misgurnus anguillicaudatus* based on SMRT sequencing, and constructed a transcriptome by performing functional annotations of the non-redundant transcripts with public databases [38]. By sequencing mixed samples of *Agasicles hygrophila* eggs, larvae, pupae, and adults using PacBio SMRT, Jia and colleagues constructed a transcripotome composed of 28,982 full-length transcripts (transcript N50 of 2331 bp) [5]. Here, we employed PacBio SMRT technology for whole-transcriptome profiling in *A. apis* mycelia. A total of ~23.97 Gb of subreads were generated, including 464,043 CCS (mean length of 2970 bp) and 394,142 FLNC reads (mean length of 2820 bp). After removing redundant sequences from 182,165 high-quality isoforms, 174,095 transcripts (mean length of 2728 bp) and 5141 genes (mean length of 4087 bp) were obtained, which is much better than the 42,609 assembled unigenes (mean length of 966 bp) recorded in our previous work [22]. Meanwhile, the non-assembled transcripts from SMRT sequencing (transcript N50 of 3543 bp) were much longer than the assembled transcripts from Illumina sequencing (unigene N50 of 1550 bp) [22]. Additionally, the expression of 16 *A. apis* isoforms was verified by RT-PCR. Among these transcripts, one was further confirmed by Sanger sequencing (**Figure 8**). The results indicated that full-length transcripts can be recovered using SMRT sequencing. Additionally, 5141 genes were detected by SMRT with a mean length of 4087 bp, which is 934 bp larger in size than that in the reference genome. About 13,853 isoforms were found to carry a completed ORF, further showing the long-read property of PacBio SMRT sequencing. The human transcriptome was revealed to be much more complex than previously expected owing to the application of TGS technology, which identified an array of novel isoforms that had not yet been annotated [9]. Similar findings were reported in some other species such as pigs [39], rabbits [26], switchgrass [13], red clover [30], and so forth. In the present study, we identified 2405 novel genic loci from unannotated regions and 11,623 new isoforms from various exons in the draft genome based on SMRT data, suggestive of a more complex transcriptome of *A. apis*. These data not only enrich the transcriptional information of the draft genome sequence but could also be used in functional studies of important genes in further research.

Here, 17,195 transcripts were annotated in four functional databases including the Nr, KOG, GO, and KEGG databases. Genomic sequencing clearly suggested that most of genes specifying the key biological functions are shared by all eukaryotes [40]. In this study, KOG database annotations showed that majority of transcripts (3,370, 31.55%) were enriched in the function of general function prediction only. The *A. apis* transcripts were annotated to various subcategories such as metabolic process, cellular process, cell, cell part, binding, and catalytic activity in the three categories based on the GO database annotations. Additionally, KEGG database annotations demonstrated that these transcripts were annotated to as many as 87 material and energy metabolism-related pathways such as biosynthesis of secondary metabolites and oxidative phosphorylation, genetic information processing-related pathways, such as RNA transport and spliceosome, cellular processes-related pathways such as cell cycle and endocytosis; environmental information processing-related pathways, such as Mitogen-activated protein kinase (MAPK) signaling pathway and ATP-binding cassette (ABC) transporters, and an organismal systems-related pathway (longevity regulating pathway). These results suggest that the transcripts of *A. apis* are associated with the abovementioned functions. Collectively, this high-quality transcriptome of *A. apis* could be used as a reference in some cases.

In a previous study, we identified 379 lncRNAs in the mixed samples of *A. apis* mycelia and spores, including 242 antisense lncRNAs, 123 intergenic lncRNAs, one intronic lncRNA and 13 sense lncRNAs, based on NGS and bioinformatics [41]. In this work, 1205 *A. apis* lncRNAs with an average length of 912 bp were identified on the basis of PacBio SMRT sequencing. These high-quality lncRNAs were much longer than previous ones [41], indicative of the advantage of SMRT sequencing in mining lncRNAs at the transcriptome scale. In addition, the *A. apis* lncRNAs identified in this study have fewer exons, shorter exon and intron lengths, shorter transcript lengths, lower expression levels, and less AS evens compared with protein-coding transcripts, which are similar to our previous findings [41]. Collectively, our data provide enrichment for the lncRNA reservoir of *A. apis*, but also enlarge the ncRNA database of the fungal kingdom. It should be noted that the occurrences of false positives of non-coding transcripts can not be absolutely excluded since this conclusion just depends on the computational approach of a homologous search against reference protein databases, thus the functions in *A. apis* require further experimental evidence.

## 4. Materials and Methods

### 4.1. Preparation of A. apis mycelia samples

*A. apis* was previously isolated from a fresh chalkbrood mummy of *A. m. ligustica* larvae [41] and kept at the Honeybee Protection Laboratory of the College of Bee Science at Fujian Agriculture and Forestry University.

*A. apis* was cultured at 33±0.5 °C on plates of Potato-Dextrose Agar (PDA) medium according to the method developed by Jensen et al. [42]. One week after culturing, mycelia (shown in **Figure 1**) were harvested and purified as previously described [42], and then immediately frozen in liquid nitrogen and stored at −80 °C.

### 4.2. Library construction and SMRT sequencing

Firstly, the total RNA was extracted by grinding *A. apis* mycelia in TRIzol reagent (Thermo Fisher, Shanghai, China) on dry ice and processed following the protocol provided by the manufacturer. The integrity of the RNA was determined with the Agilent 2100 Bioanalyzer and agarose gel electrophoresis. The purity and concentration of the RNA were determined with the Nanodrop micro-spectrophotometer (Thermo Fisher, Shanghai, China). Secondly, mRNA was enriched by Oligo (dT) magnetic beads, followed by reverse transcription of the enriched mRNA into cDNA using Clontech SMARTer PCR cDNA Synthesis Kit (Takara, Shiga, Japan). PCR cycle optimization was used to determine the optimal amplification cycle number for the downstream large-scale PCR reactions. Then the optimized cycle number was used to generate double-stranded cDNA. Thirdly, >4kb size selection was performed using the BluePippin™ Size-Selection System (Select science, Corston, UK) and mixed equally with the no-size-selection cDNA. Fourthly, large-scale PCR was performed for the next SMRT bell library construction; cDNAs were DNA damage repaired, end repaired, and ligated to sequencing adapters. Finally, the SMRT bell template was annealed to sequencing primer and bound to polymerase followed by sequencing on the PacBio Sequel platform using P6-C4 chemistry with 10 h movies by Gene Denovo Biotechnology Co. (Guangzhou, China).

### 4.3. Illumina short-read sequencing

(1) The total RNA was isolated from *A. apis* mycelia using a Trizol Kit (Thermo Fisher, Shanghai, China). (2) Oligo (dT) primers were used to isolate poly-A mRNA, followed by fragmentation and reverse transcription with random primers (Qiagen, Hilden, Germany). Second-strand cDNAs were synthesized using RNase H and DNA polymerase I. The double-strand cDNAs were then purified using the QiaQuick PCR extraction kit (Qiagen, Hilden, Germany). (3) After agarose gel electrophoresis, the required fragments were purified using a DNA extraction kit (Qiagen, Hilden, Germany) and then enriched via PCR amplification in a total volume of 50 μL containing 3 μL of NEB Next USER Enzyme (NEB, Ipswich, USA), 25 μL of NEB Next High-Fidelity PCR Master Mix (2×) (NEB, Ipswich, USA), 1 μL of Universal PCR Primer (25 mmol) (NEB, Ipswich, USA), and 1 μL of Index (X) Primer (25 mmol) (NEB, Ipswich, USA). The reaction conditions were as follows: 98 °C for 30 s, followed by 13 cycles of 98 °C for 10 s and 65 °C for 75 s, and 65 °C for 5 s. (4) The amplified fragments were sequenced on the Illumina HiSeq™ 4000 platform (Illumina, San Diego, USA) by Gene Denovo Biotechnology Co. (Guangzhou, China) following the manufacturer’s protocols.

### 4.4. Processing of SMRT reads and error correction

The raw sequencing reads of cDNA libraries were classified and clustered into transcript consensus using the SMRT Link v5.0.1 pipeline [43] supported by Pacific Biosciences. Briefly, CCS reads were extracted out of subreads of the BAM file with a minimum full pass of 1 and a minimum read score of 0.65. Subsequently, CCS reads were classified into FLNC, non-full-length (nFL), chimeras, and short reads based on cDNA primers and the poly-A tail signal. Reads shorter than 50 bp were discarded. Next, the FLNC reads were clustered by ICE software to generate the cluster consensus isoforms.

Two strategies were employed to improve the accuracy of PacBio reads: (1) The nFL reads were used to polish the above obtained cluster consensus isoforms by Quiver software to obtain the FL-polished high-quality consensus sequences (accuracy≯99%). (2) The low-quality isoforms were further corrected using Illumina short reads obtained from the same samples using the LoRDEC tool (version 0.8) [44]. The pipeline of the SMRT sequencing data process is shown in **Figure 1**.

### 4.5. Mapping of PacBio data to reference genome

The corrected high quality consensus sequences were then mapped to the reference genome of *A. apis* (AAP 1.0) using Genomic Mapping and Alignment Program (GMAP) [45], and redundant transcripts were collapsed with minimum identity of 95% and a minimum coverage of 99%. The finally obtained isoforms were compared with the reference genome annotation and classified into three groups: known isoforms, novel isoforms, and new isoforms.

### 4.6. Functional annotation of transcripts

Transcripts were aligned against the NCBI Nr database (http://www.ncbi.nlm.nih.gov), KOG database (http://www.ncbi.nlm.nih.gov/KOG), and KEGG database (http://www.genome.jp/kegg) with the BLASTx program (http://www.ncbi.nlm.nih.gov/BLAST/) at an E-value threshold of 1×10^-5^ to evaluate the sequence similarity with genes of other species. GO annotation was analyzed by Blast2GO software [46] with the Nr annotation results of isoforms. Isoforms ranking as having the highest 20 scores and no shorter than 33 High-scoring Segment Pair (HSP) hits were selected for the Blast2GO analysis. Functional classification of isoforms was then performed using WEGO software [47].

### 4.7. ORF prediction

The ORFs were detected by using ANGEL [48] software for the isoform sequences to obtain the coding sequences (CDS), protein sequences, and (Untranslated region) UTR sequences.

### 4.8. Prediction and analysis of lncRNAs

CNCI (version 2) [49] and CPC [49] (http://cpc.cbi.pku.edu.cn/) were used to evaluate the protein-coding potential of novel isoforms and new isoforms by default parameters. Meanwhile, isoforms were mapped to the SwissProt database to assess protein annotation. The intersection of both non protein-coding potential results and non-protein annotation results were regarded as candidate lncRNAs. To better annotate lncRNAs at the evolution level, Infernal [50] (http://eddylab.org/infernal/□ was used to assess the secondary structures and sequence conservation of lncRNAs. Cuffcompare was used to select the different types of lncRNAs including lincRNA, intronic lncRNA, and anti-sense lncRNA. Fragments per kilobase per million fragments mapped (FPKM) of both lncRNAs and mRNAs were calculated using StringTie (1.3.1). The transcript lengths, exon numbers and lengths, intron lengths, GC content, expression levels, and AS event numbers of lncRNAs were compared with those of mRNAs.

### 4.9. TF analysis

Protein coding sequences of isoforms were aligned by hmmscan to Plant TFdb (http://planttfdb.cbi.pku.edu.cn/) to predict TF families.

### 4.10. RT-PCR validation of isoforms

Sixteen isoforms of *A. apis* (Isoform000014, Isoform000018, Isoform000019, Isoform000021, Isoform000027, Isoform000028, Isoform000029, Isoform000042, Isoform000047, Isoform000063, Isoform000066, Isoform000085, Isoform000094, Isoform000113, Isoform000127 and Isoform0000365) were randomly selected for RT-PCR validation. Specific forward and reverse primers (presented in Table S1) were designed using DNAMAN software on the basis of the corresponding transcript sequences. One microgram of total RNA of *A. apis* mycelia was reverse transcribed to cDNA using the RevertAid First Strand cDNA Synthesis Kit (TaKaRa, China) and Oligo dT primers. PCR amplification was conducted on a T100 thermo cycler (BIO-RAD) using Premix (TaKaRa, China) under the following conditions: pre-denaturation step at 94 °C for 5 min; 34 amplification cycles of denaturation at 94 °C for 30 s, annealing at 60 °C for 30 s, and elongation at 72 °C for 1 min; this was followed by a final elongation step at 72 °C for 10 min. The PCR products were monitored on 1.8% agarose gel through electrophoresis with Genecolor (Gene-Bio, China) staining. The fragment amplified from Isoform000014 was purified and cloned to pMD-19T vector (Takara, China) followed by Sanger sequencing.

### 4.11. Data availability

The raw data produced from PacBio SMRT sequencing and Illumina sequencing in this work were submitted to NCBI SRA database under BioProject numbers: PRJNA557811 and PRJNA560452.

## 5. Conclusions

Taken together, this work, for the first time, proposed the full-length transcriptome of *A. apis* using PacBio SMRT sequencing, providing a basis for further exploration of gene structures such as AS and APA. Moreover, the annotation of the *A. apis* gene set could improve the reference genome annotation and facilitate deeper understanding of the complexity of the *A. apis* genome and transcriptome.

## Author Contributions

Conceptualization, D.C. and R.G. designed this study. R.G., Y.D., X.F., Z.Z., J.W., H.J., Y.F., H.C., D.Z., X.X., Q.L., C.X. and Y.Z. conducted laboratory work. R.G. and Y.D. performed bioinformatic analysis. D.C., R.G. and Y.D. supervised the work and contributed to preparation of the manuscript. All authors read and approved the final manuscript.

## Funding

This research was supported by the Earmarked Fund for China Agriculture Research System (No. CARS-44-KXJ7), the Science and Technology Planning Project of Fujian Province (No. 2018J05042), the Teaching and Scientific Research Fund of Education Department of Fujian Province (No. JAT170158), the Outstanding Scientific Research Manpower Fund of Fujian Agriculture and Forestry University (No. xjq201814), and the Scientific and Technical Innovation Fund of Fujian Agriculture and Forestry University (No. CXZX2017342, No. CXZX2017343).

## Acknowledgments

We thank all editors and reviewers for their invaluable comments.

## Conflicts of Interest

The authors declare no conflict of interest.

## Abbreviations

ABC: ATP-binding cassette
APA: Alternative polyadenylation
AS: Alternative splicing
CCS: Circular consensus sequence
CDS: Coding sequences
CNCI: Coding-Non-Coding Index
CPC: Coding potential calculator
FLNC: Full-length non-chimeric
FPKM: Fragments per kilobase per million fragments mapped
GMAP: Genomic Mapping and Alignment Program
GO: Gene Ontology
HSP: High-scoring Segment Pair
ICE: Iterative Clustering for Error Correction
KEGG: Kyoto Encyclopedia of Genes and Genomes
KOG: Clusters of euKaryotic Orthologous Groups
LncRNA: Long non-coding RNA
MAPK: Mitogen-activated protein kinase
nFL: Non-full-length
NGS: Next-generation sequencing
Nr: NCBI non-redundant protein
ORF: Open reading frame
PDA: Potato-Dextrose Agar
SMRT: Single molecule real time
TF: Transcription factor
TGS: Third-generation sequencing
UTR: Untranslated region

---

**Table S1.**
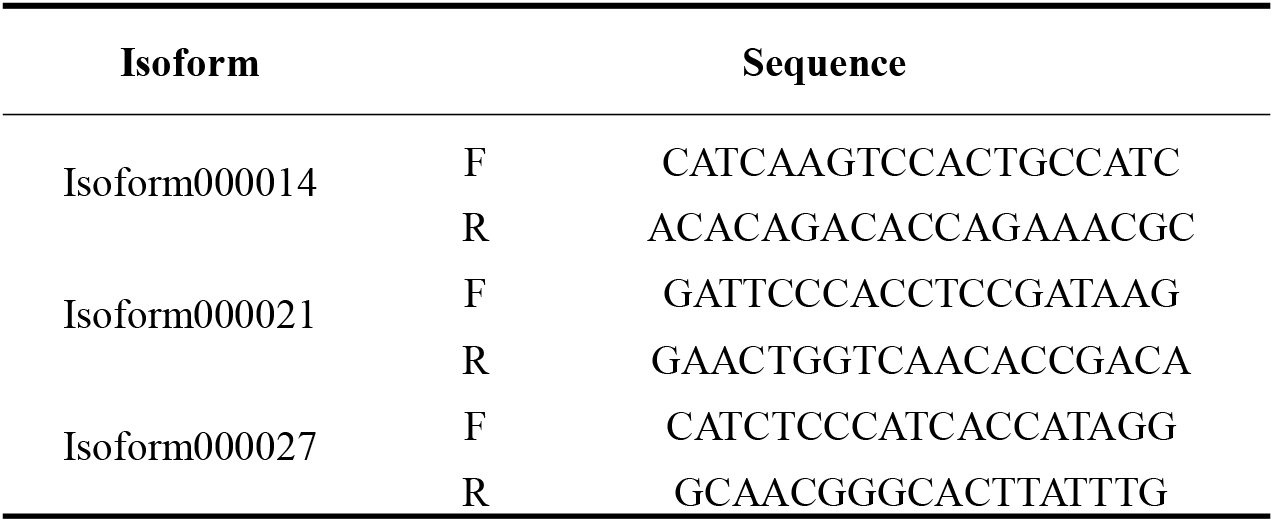
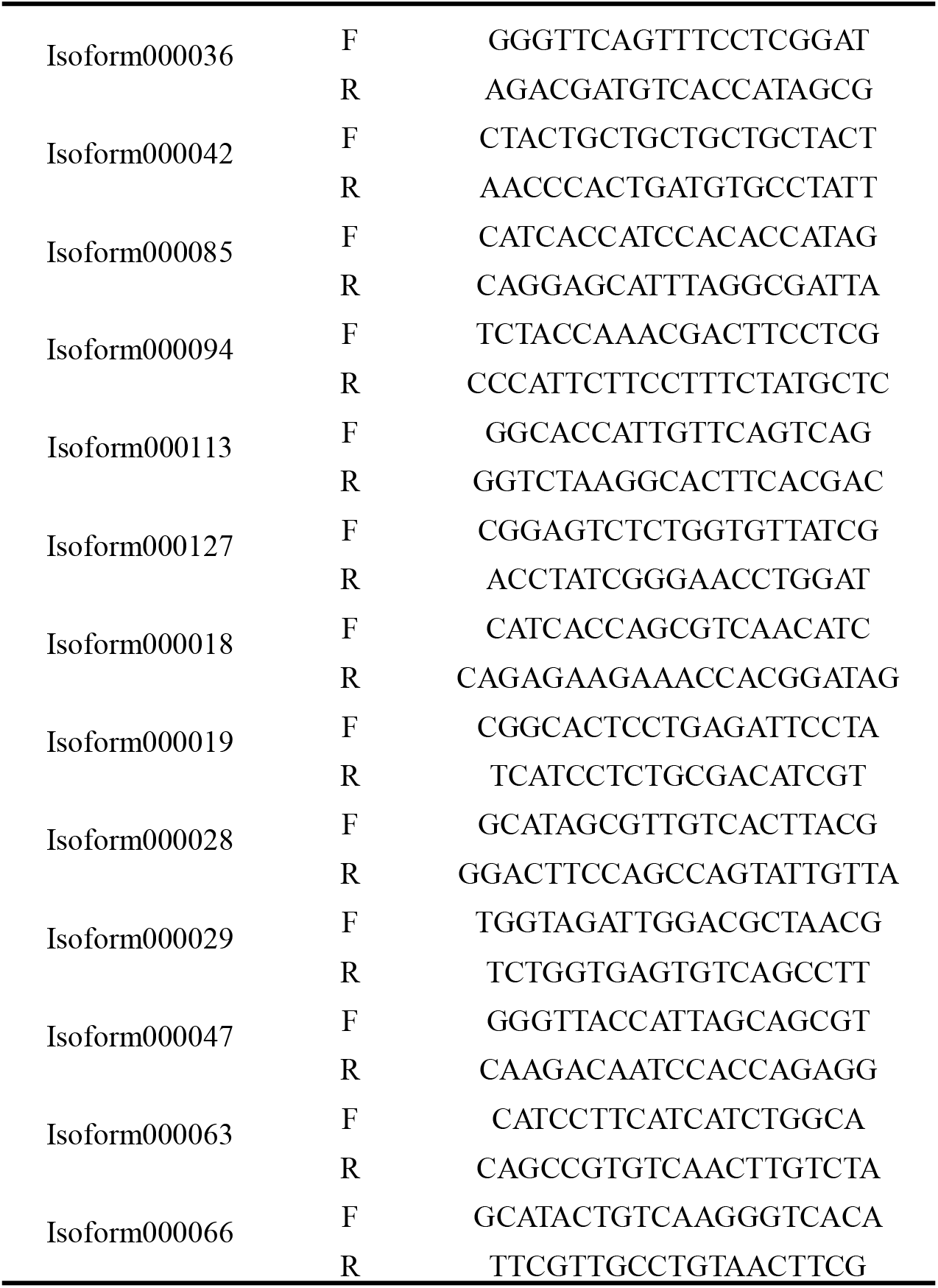
Primers used in the present study.

